# The evolution of mutual mate choice under direct benefits

**DOI:** 10.1101/046714

**Authors:** Alexandre Courtiol, Loïc Etienne, Romain Feron, Bernard Godelle, François Rousset

## Abstract

In nature, the intensity of mate choice (*i.e*., choosiness) is highly variable within and between sexes. Despite growing empirical evidence showing male and/or mutual mate choice, theoretical investigations of the joint evolution of female and male choosiness are few. In addition, previous approaches have often assumed an absence of trade-off between the direct benefits per mating and the lower mating rate that results from being choosy. Here, we model the joint evolution of female and male choosiness when it is solely ruled by this fundamental trade-off. We show that this trade-off can generate a diversity of stable combinations of choosiness. Mutual mate choice can only evolve if both females and males exhibit long latency after mating. Further, we show that an increase in choosiness in one sex does not necessarily prevent the evolution of mutual mate choice: the outcome depends on details of the life history, the decision rule for mate choice, and how the fecundity of a pair is shaped by the quality of both individuals. Lastly, we discuss the power of the sensitivity of the relative searching time (*i.e*., of the proportion of lifetime spent searching for mates) as a predictor of the joint evolution of choosiness.

## Introduction

Mate choice corresponds to any behavior that increases (or decreases) the probability of mating with certain individuals (Halliday, 1983). Darwin (1871) proposed mate choice as the mechanism responsible for the evolution of extravagant ornaments. Because males generally display these ornaments, the first empirical investigations of mate choice were highly focused on females. However, recent research has shown that the intensity of choice (*i.e*., choosiness) varies widely across taxa both within and between sexes. In particular, empirical evidence for male mate choice keeps accumulating (for reviews, see Clutton-Brock, 2009; Edward and Chapman, 2011). Besides, mutual mate choice – the situation in which both females and males are choosy – has also been documented in a wide variety of taxonomic groups, including amphibians (Verrell, 1995), arachnids (Rypstra et al., 2003; Cross et al., 2007; Luo et al., 2014), birds (Jones and Hunter, 1993; Monaghan et al., 1996; Hansen et al., 1999; Faivre et al., 2001; Sæther et al., 2001; Romero-Pujante et al., 2002; Daunt et al., 2003; Pryke and Griffith, 2007; Nolan et al., 2010), crustaceans (Aquiloni and Gherardi, 2008), fishes (Rowland, 1982; Hua Wen, 1993; Kraak and Bakker, 1998; Sandvik et al., 2000; Werner and Lotem, 2003; Wong et al., 2004; Bahr et al., 2012; Myhre et al., 2012), insects (for a review, see Bonduriansky, 2001) and mammals (Drickamer et al., 2003) including primates (Gomez et al., 2012; Courtiol et al., 2010). Despite the ever-growing empirical literature showing that choosiness is highly variable in both sexes, theoretical investigations of the joint evolution of female and male choosiness are few compared to the large number of studies dealing with unilateral mate choice (Bergstrom and Real, 2000).

Why does choosiness vary so much both within and between sexes and species? One potential explanation is that its evolution is influenced by benefits and costs that vary due to differences in life-history traits and/or environmental conditions (Jennions and Petrie, 1997). Mate choice is indeed often associated with direct fitness benefits (*e.g*., nuptial gifts, territory, food, protection, increased fertility, or parental care; Andersson, 1994) and costs (*e.g*., increased predation risk or injuries caused by conspecifics; Andersson, 1994) for the chooser, regardless of its sex. The presence of these direct benefits and costs in a wide variety of organisms suggests that direct selection plays an important role in the evolution of mate choice (Jones and Ratterman, 2009). However, predicting variation in the direct selection of choosiness is difficult because the nature of benefits and costs involved often depends on the organism being studied.

One general cost that has been ignored by most theoretical studies in sexual selection is that choosy individuals necessarily suffer a decrease in their mating rate (Etienne et al., 2014; Dechaume-Moncharmont et al., 2016). This is because choosy individuals spend time searching for particular mates instead of reproducing with the first member of the other sex they encounter. When mating events are sequential, the total number of matings for choosy individuals is thus reduced and this cost – sometimes qualified as an *opportunity* cost – occurs even if individuals only mate once in their lives (because rejecting mates increases the probability of dying before having reproduced). Mate choice is thus intrinsically associated with a trade-off between the benefits per mating and the mating rate that, respectively, increase and decreases with choosiness (Owens and Thompson, 1994; Kokko and Mappes, 2005; Härdling et al., 2008). We call this *the fundamental trade-off of mate choice*. Etienne et al. (2014) evaluated the importance of this trade-off under the assumption that mate choice can only evolve in one sex (the other being considered indiscriminate). They showed that, depending on the biological and ecological context, the strength of the trade-off varies and influences the evolution of choosiness.

Here, we extend the model of Etienne et al. (2014) to study the influence of this fundamental trade-off of mate choice when choosiness is allowed to evolve in both sexes. Such a generalization is not trivial because the evolution of choosiness in one sex influences the evolution of choosiness in the other (Johnstone et al., 1996; Johnstone, 1997; Kokko and Johnstone, 2002). Indeed, choosiness in each sex impacts on the competition for mates in the other sex, which influences in turn the benefits and costs associated with choosiness in both sexes. Or as Johnstone (1997) put it: “the best strategy for males depends on the behaviour of females, and *vice versa*”. We also attempt to obtain a simple metric that allows for general predictions about the evolution of choosiness, when the trade-off is the sole evolutionary force shaping mate choice. Etienne et al. (2014) showed that, within this scope, the evolution of choosiness in one sex can be predicted in terms of the proportion of a lifetime devoted to searching for mates or RST for short (*i.e*., the Relative Searching Time). More specifically, the sensitivity of RST (*i.e*., *∂*RST) – the change in RST caused only by a variation in any biological or ecological parameter affecting the mating rate of individuals, while choosiness is fixed – gives the effect of such variation on selection on choosiness. When *∂*RST is positive, lower choosiness is selected, and *vice versa*. Here, we investigate the predictive power of *∂*RST on the joint evolution of female and male choosiness.

Factors other than the fundamental trade-off of mate choice certainly influence the evolution of choosiness (*e.g*., indirect benefits, sexual conflicts). Yet, we chose to study the influence of this trade-off in isolation for two main reasons. First, the evolutionary consequences of this trade-off have been shown to be complex even when choosiness is free to evolve in only one sex (Etienne et al., 2014). Second, these other sources of selection are likely to act in addition to, and not instead of, the trade-off we consider. Hence, computing how this trade-off influences the direct selection of choosiness should help to disentangle the impacts of the various different selection pressures that shape the evolution of mate choice. In particular, we consider the evolution of choosiness given a previously established pattern of parental investment in each sex and do not study the joint evolution that could occur between choosiness and parental care (Kokko and Jennions, 2008). This assumption allows us to study the fundamental trade-off of mate choice independently from the one between mating rate and parental care.

Our work complements existing theoretical studies on the evolution of mutual mate choice. Specifically, we consider a continuous strategy set for choosiness and thereby extend previous studies which considered two discrete categories of choosiness (Crowley et al., 1991; Härdling et al., 2008). Moreover, our formalism relies on a game-theoretic approach allowing a full consideration of the influence of other-sex and same-sex individuals’ behaviors on the evolution of choosiness, contrary to some earlier models (Owens and Thompson, 1994; Kokko and Monaghan, 2001; Simao and Todd, 2002; Kokko and Mappes, 2005; Gowaty and Hubbell, 2009). As such, our approach complements the study by Johnstone et al. (1996) that focused on the diversity of mating patterns emerging from mutual mate choice, and the one by Kokko and Johnstone (2002) that focused on the joint evolution between choosiness, signaling and care. Finally, we allow for any number of mating events throughout a lifetime, generalizing models that assume that individuals mate only once (Parker, 1983; McNamara and Collins, 1990; Johnstone, 1997; Alpern and Reyniers, 1999; Alpern and Reyniers, 2005; Alpern and Katrantzi, 2009; Ramsey, 2011).

## The model

### Individual traits

We consider an infinite population at demographic equilibrium with two sexes in equal proportion (sex-ratio = 1:1). One sex, denoted *x*, is treated as the focal sex. The other is denoted *y*. Each individual *i* of sex *x* is characterized by a quality *q_x_*_,*i*_ and a choosiness *ϕ_x_*_,*i*_, both real numbers between 0 and 1 (see Table 1 for a summary of our notations). We assume *q_x_*_,*i*_ to be directly proportional to the contribution of a mate to the fecundity of a given mating (*i.e*., it directly translates into offspring quantity), which is why we call it *quality*. Quality is strictly environmentally determined and follows a beta distribution (with its two parameters denoted *α_x_* and *β_x_*) which we assume to be constant across generations. This assumption prevents the emergence of linkage disequilibrium between choosiness and quality. Thus, there is no indirect selection of choosiness (and thereby no so-called *good genes*) in this model. Choosiness sets the proportion of other-sex individuals whose quality is too low to be accepted as mates. For instance, an individual *i* of sex *x* with *ϕ_x_*_,*i*_ = 0.4 rejects all individuals of sex *y* in the lower 40% of the quality distribution and accepts all those whose quality is higher. We denote *q_y_*(*ϕ_x_*_,*i*_) the minimal value of quality in sex *y* that is accepted by the individual *i* of sex *x*. Thus, *q_x_* and *q_y_* correspond to quantile functions in each sex. We assume that individuals make no error in assessing the quality of their potential mates and that this trait is strictly genetically determined by one sex-specific locus and is expressed as a fixed threshold. Choosiness is therefore considered to be independent from individual qualities.

**Table 1:**
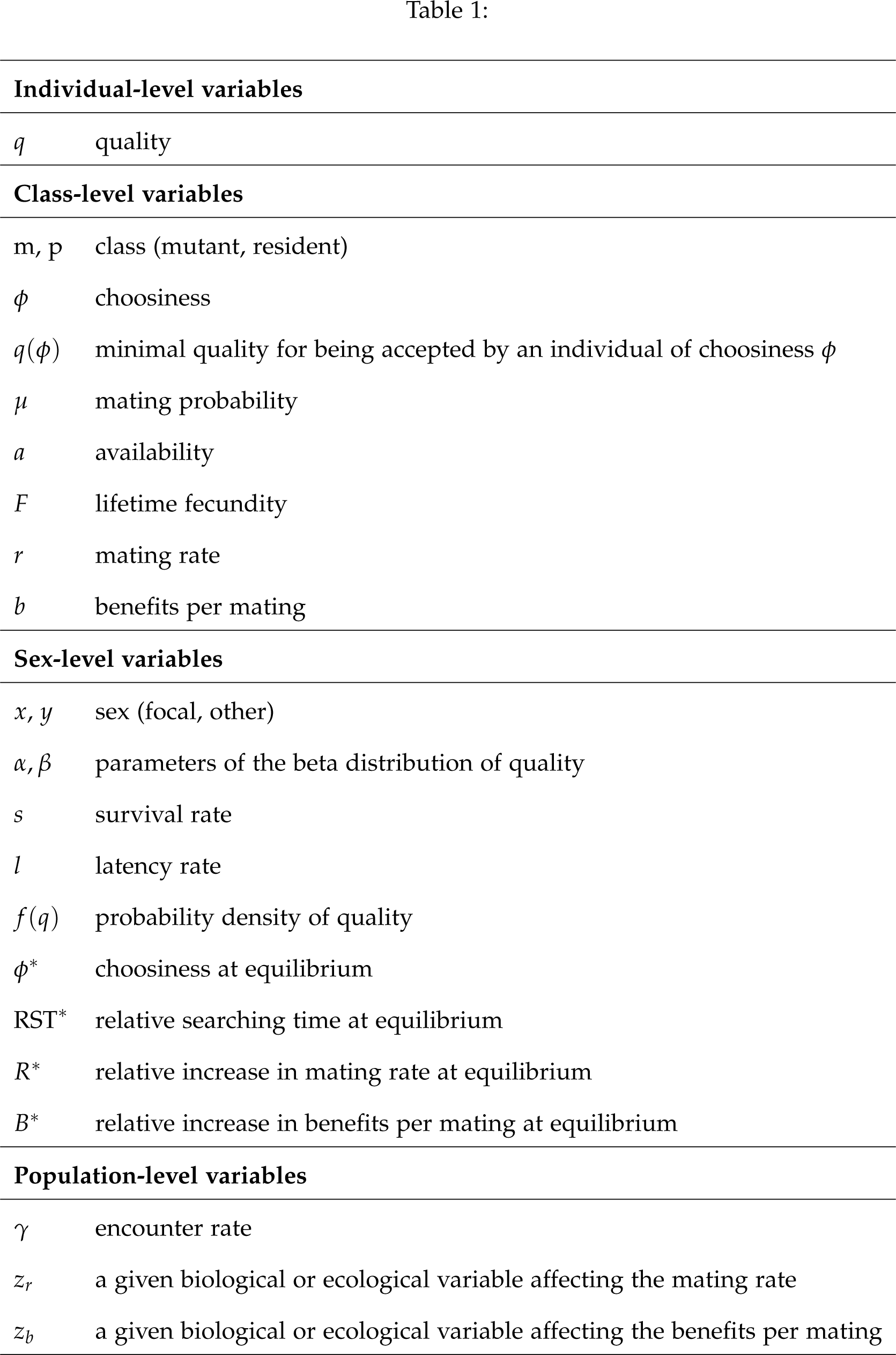
Summary of notation.

|  |  |
| --- | --- |
| **Individual-level variables** |  |
| $q$ | quality |
| **Class-level variables** |  |
| $m, p$ | class (mutant, resident) |
| $\phi$ | choosiness |
| $q(\phi)$ | minimal quality for being accepted by an individual of choosiness $\phi$ |
| $\mu$ | mating probability |
| $a$ | availability |
| $F$ | lifetime fecundity |
| $r$ | mating rate |
| $b$ | benefits per mating |
| **Sex-level variables** |  |
| $x, y$ | sex (focal, other) |
| $\alpha, \beta$ | parameters of the beta distribution of quality |
| $s$ | survival rate |
| $l$ | latency rate |
| $f(q)$ | probability density of quality |
| $\phi^*$ | choosiness at equilibrium |
| $RST^*$ | relative searching time at equilibrium |
| $R^*$ | relative increase in mating rate at equilibrium |
| $B^*$ | relative increase in benefits per mating at equilibrium |
| **Population-level variables** |  |
| $\gamma$ | encounter rate |
| $z_r$ | a given biological or ecological variable affecting the mating rate |
| $z_b$ | a given biological or ecological variable affecting the benefits per mating |

### The life cycle

Time is discrete, and at each time step, each individual of sex *x* survives with rate *s_x_*. We consider that *s_x_* is independent from all other individual traits (*i.e*., *q_x_*_,*i*_ and *ϕ_x_*_,*i*_ have no effect on survival). The expected lifetime of individuals of sex *x* is thus 1/(1 *− s_x_*) (the time step during which an individual dies is included in its lifetime). At any time step for which an individual of sex *x* survives, it randomly encounters an individual of sex *y* with rate *γ*. (Due to the balanced sex-ratio, individuals of sex *y* also encounter individuals of sex *x* at the same rate *γ*.) If both individuals are available and accept each other, mating occurs. In this case, mated individuals of sex *x* enter a latency period with rate *l_x_* during which they become unavailable for mating. Biologically, latency can result from any process that prevents individuals from remating instantly (gamete depletion, mate guarding, parental care, etc.). Then, at each time step, the latent individual, if it survives, remains in latency with rate *l_x_*. We therefore assume that the duration of latency is independent between the male and the female of a given mating pair. Once latency is finished, the individual becomes available for mating again. The transition rates between “available” and “unavailable” states are thus given by the following matrix (see also Etienne et al., 2014):

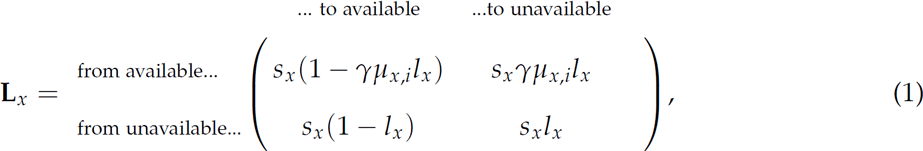

where *µ_x_*_,*i*_ is the probability that an individual *i* of the sex *x* mates, given that it is available for mating and has encountered an individual of sex *y*. Similarly, the transition rates for individuals of sex *y* are obtained by substituting *x* for *y* in the previous matrix.

### Calculating mating probabilities

The mating probability *µ_x_*_,*i*_ of an individual *i* of sex *x* depends on the probability that it finds the potential mate encountered acceptable, thus on its choosiness (*i.e*., *ϕ_x_*_,*i*_). In addition, *µ_x_*_,*i*_ depends on the availability of individuals of sex *y*, that is on the probability that a given individual of sex *y* is not in latency. This availability is in turn related to the choosiness of other individuals of sex *x*. The reason is that an individual that is encountered may be in latency after having previously mated – and is thus unavailable for a new mating. To take this competition for mates into account, we consider a mutant individual *m* with choosiness *ϕ_x_*_,m_ in a population where all other individuals of sex *x* have choosiness *ϕ_x_*_,p_ (with *p* for population). We also assume that all individuals of sex *y* show the same choosiness, denoted *ϕ_y_*_,p_. Together, *ϕ_x_*_,p_ and *ϕ_y_*_,p_ define the residents in the population.

We now characterize the relationship between the mating probability *µ_x_*_,m_ of a mutant of the focal sex and the other parameters. First, *µ_x_*_,m_ depends on the quality of the mutant (*i.e. q_x_*_,m_). Indeed, if the latter is not of sufficient quality to mate with (*i.e*., with quality *q_x_*_,m_ < *q_x_*(*ϕ_y_*_,p_)), it is never chosen by other-sex individuals and thus its mating probability is null. If so, it does not transmit its choosiness alleles and thus does not influence the evolution of choosiness in the population. Therefore, only mutants who can obtain mates need to be taken into account. Two situations need to be distinguished for such a mutant. First, if it is choosier than other same-sex individuals (*ϕ_x_*_,m_ ≥ *ϕ_x_*_,p_), the potential partners it is willing to mate with are also courted by residents and are thus not necessarily available. The availability of such potential partners, that is the probability that any individual *i* of sex *y*, with quality *q_y_*_,*i*_ ≥ *q_y_*(*ϕ_x_*_,p_), is in the available state in eq. 1, is denoted *a_y_*_,p_. Second, if the mutant is less choosy than residents (*ϕ_x_*_,m_ < *ϕ_x_*_,p_), it is willing to mate with two types of individuals: those who are also chosen by resident individuals of sex *x*, whose availability equals *a_y_*_,p_, and those whose quality ranges from *q_y_*(*ϕ_x_*_,m_) to *q_y_*(*ϕ_x_*_,p_) and who are thus always available for mating with this mutant. Therefore, we have:

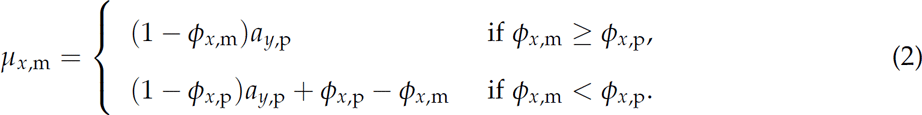

To characterize the mating probability *µ_x_*_,p_ of a focal-sex resident whose quality is sufficient to mate with (*i.e*., with quality *q_x_*_,*i*_ ≥ *q*_*x*_(*ϕ_y_*_,p_)), we set *ϕ_x_*_,m_ = *ϕ_x_*_,p_ in the previous equation. We obtain:

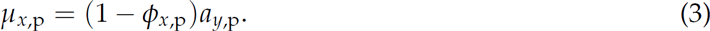

### Calculating mating availabilities

To obtain the expressions for the mating availability *a_y_*_,p_, we need to compute, in each sex, the expected time spent by resident individuals in latency and to divide it by the expected lifespan. Because the states of the life cycle considered here forms a Markov chain where death is an absorbing state, the expected time spent in each state can be deduced from the transition probabilities between the non-absorbing states of the life cycle (using **D**_*x*_ = (**I** *−* **L**_*x*_)^−1^ with **I** the identity matrix and **L**_*x*_ from eq. 1, see *e.g*., Caswell, 2001, p. 112). Assuming that individuals start their reproductive life available for mating, we can therefore deduce the average number of time steps *d* (first element of the matrix **D**_*x*_) that a focal-sex resident spends available for mating throughout lifetime:

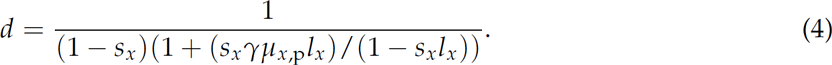

By dividing *d* by the expected lifespan (1/(1 *− s_x_*)) and substituting *µ_x_*_,p_ for the value obtained from eq. 3, we obtain the probability *a_x_*_,p_ which represents the availability of residents of sex *x* whose quality is sufficient to mate (*i.e*., with quality *q_x_*_,*i*_ ≥ *q*_*x*_(*ϕ_y_*_,p_)). Substituting *x* for *y*, we similarly obtain the availability *a_y_*_,p_ for a resident of sex *y* whose quality is sufficient to mate (*i.e*., with quality *q_y_*_,*i*_ ≥ *q*_*y*_(*ϕ_x_*_,p_)) at a given time step. This leads to the following system of two equations with two unknowns:

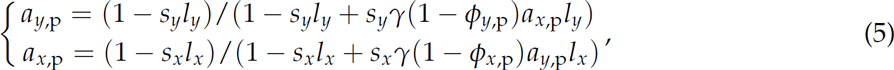

which solution yields:

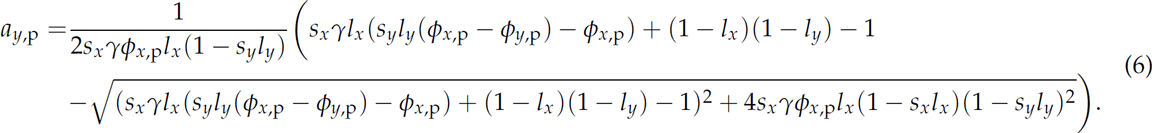

Exchanging *x* and *y* in this expression gives *a_x_*_,p_.

As further computations require the expression of the availability of a mutant *m* of sex *x*, we used the same approach to compute *a_x_*_,m_ and obtained:

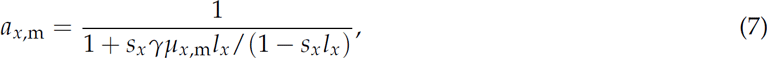

where *µ_x_*_,m_ (that is a function of *a_y_*_,p_) is given by eq. 2.

### Computing the expected lifetime fecundity of a mutant

Let us define the lifetime fecundity of an individual *i* as the number of offspring it produces as a result of all mating events. We define the *expected* lifetime fecundity as the lifetime fecundity computed in a lineage of individuals. That is, the expected lifetime fecundity is computed over the distribution of contexts in which an individual of this lineage could be. To obtain this expected lifetime fecundity, we first compute the expected fecundity *F_x_*(*q_x_*_,*i*_) of an individual *i* of sex *x* given its quality *q_x_*_,*i*_. Then, we will compute its expectation over the distribution of quality of *q_x_*_,*i*_. For these computations, we assume that the number of offspring obtained from any mating (*i.e*., the benefits per mating) depends neither on the number of previous matings nor on the number of offspring obtained from these previous matings. Therefore, by Wald’s formula for optional stopping (*e.g*., Durrett, 2010, p. 185), *F_x_*(*q_x_*_,*i*_) is the product of the individual’s mating rate (*r_x_*_,*i*_), its expected benefits per mating (integrated over the distribution of each partner’s quality) which we call *b*(*q_x_*_,*i*_), and its expected lifetime (1/(1 *− s_x_*)):

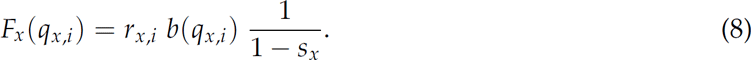

To compute the expected benefits per mating *b*(*q_x_*_,*i*_), we assume the reproductive success of a mating pair to be equal to the mean of qualities of the two members of the pair, which makes it linear in the individual quality *q_x_*_,*i*_ and in the expected quality 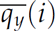 of its mates:

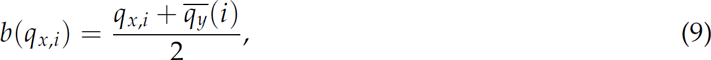

The fact that all individuals of sex *y* are assumed to have the same choosiness (see above) implies that among individuals with different *q_x_*_,*i*_ above the threshold of sex *y*, *r_x_*_,*i*_ is independent of *q_x_*_,*i*_, and individuals of lower quality never mate. Further, *q_y_*(*i*) differs among individuals with different choosiness but is identical among individuals with the same choosiness. Thus the expected lifetime fecundity *F_x_*_,m_ among all mutants representing a mutant lineage can be written as the product of expected values of the different terms of *F_x_*(*q_x_*_,*i*_):

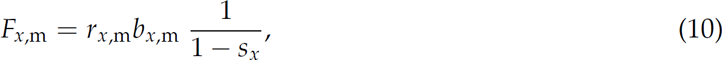

in terms of expected mating rate *r_x_*_,m_, and expected benefits per mating *b_x_*_,m_, of mutants. We will now detail expressions for these expectations.

The expected mating rate *r_x_*_,m_ of a focal-sex mutant equals its availability (*a_x_*_,m_) multiplied by the probability that it finds an individual of the other sex and mates with it at this time step (*s_x_γµ_x_*_,m_). From eq. 7, this is:

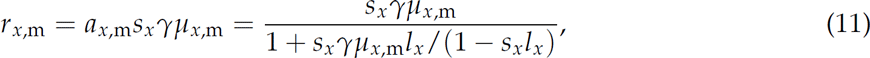

From the expression for *µ_x_*_,m_ (eq. 2), this becomes:

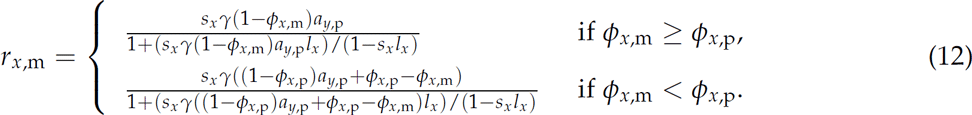

The expected benefits per mating of a mutant is the mean of the respective terms in eq. 9, which we write 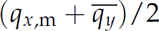. Because other-sex resident individuals accept any focal-sex individual whose quality is higher than *q_x_*(*ϕ_y_*_,p_), the expected quality of the mutant *q_x_*_,m_ is the mean of the quality distribution in sex *x* restricted to the range between *q_x_*(*ϕ_y_*_,p_) and 1. This can be written:

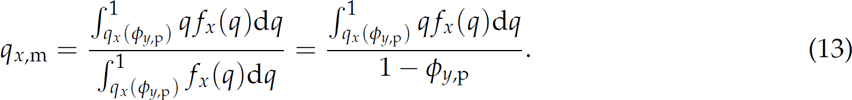

where *f_x_*(*q*) denotes the probability density of quality in sex *x*, and where the denominator of the right-hand side results from the definition of *ϕ_y_*_,p_ as the proportion of other-sex individuals whose quality is too low to be accepted as mates.

We need to distinguish two cases when computing the expected quality of the mutant’s mate (*q_y_*). First, if the mutant is choosier than resident individuals of its sex (*ϕ_x_*_,m_ ≥ *ϕ_x_*_,p_), it accepts any individual of sex *y* whose quality is higher than *q_y_*(*ϕ_x_*_,m_). In this case, the expected quality of its mates is thus the mean of the quality distribution in sex *y* restricted to the range between *q_y_*(*ϕ_x_*_,m_) and 1. Second, if the mutant is less choosy than resident individuals of its sex (*ϕ_x_*_,m_ < *ϕ_x_*_,p_), it can mate with two types of individuals who differ in their availabilities: those whose quality ranges from *q_y_*(*ϕ_x_*_,m_) to *q_y_*(*ϕ_x_*_,p_) (who are always available) and those whose quality is higher than *q_y_*(*ϕ_x_*_,p_) (who are also courted by focal-sex resident individuals and thus are available with probability *a_y_*_,p_). In this case, the expected quality of the mates of the mutant is thus the mean of the quality distribution in the sex *y* restricted to the range between *q_y_*(*ϕ_x_*_,m_) and 1, weighted by the respective availabilities of the two kinds of potential mates. By denoting *f_y_*(*q*) the density of the distribution of quality in sex *y*, we therefore have:

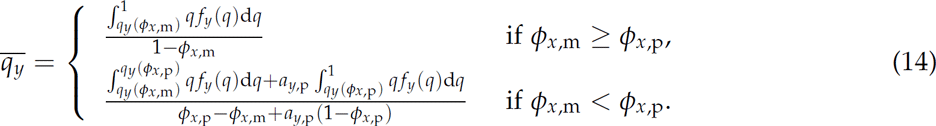

The general expression for the expected benefits per mating of a mutant is the average of the expressions for *q_x_*_,m_ and 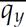:

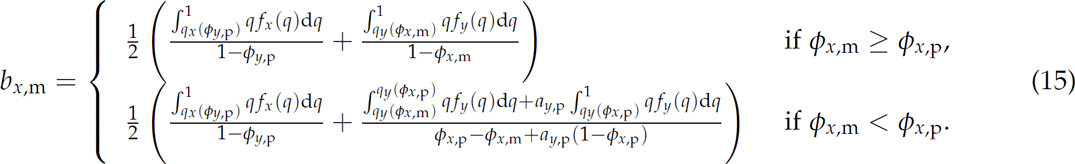

In some particular cases, the expected benefits per mating of a mutant (*b_x_*_,m_) take a simple form. For example, if the mutant is choosier than the resident (*i.e*., *ϕ_x_*_,m_ ≥ *ϕ_x_*_,p_) and if quality is uniformly distributed in both sexes (*i.e*., *f_x_*(*q*) and *f_y_*(*q*) are the beta distribution with *α_x_* = *β_x_* = *α_y_* = *β_y_* = 1), then the expected quality of the focal-sex mutant lineage and of mates are respectively (1 + *ϕ_y_*_,p_)/2 and (1 + *ϕ_x_*_,m_)/2 (as *q*(*ϕ*) = *ϕ* under the uniform distribution). In this case, the expected benefits per mating of the mutant is simply given by:

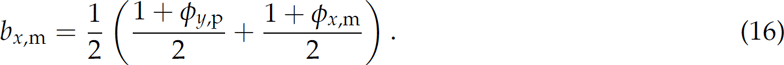

### Analytical study of the model

The full analytical methods are described in Supplementary Information, but all key steps will be presented here. We first assessed the existence of a joint equilibrium for choosiness (*i.e*., a situation in which both sexes are simultaneously at an equilibrium for choosiness) and studied its convergence and evolutionary stability (*sensu* Eshel, 1996) using standard methods from adaptive dynamics (Metz et al., 1996; Rousset, 2004). A joint equilibrium, if it exists, corresponds to the joint solution 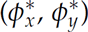 of the following system:

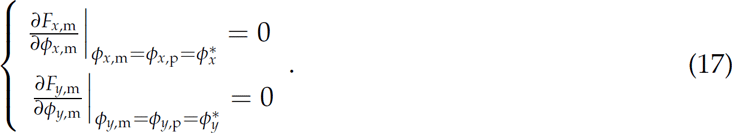

We identified such a solution and studied the convergence stability in each sex before investigating the joint convergence stability. The study of the joint convergence stability required the additional assumption of independent mutational effects between females and males. We also assessed the evolutionary stability in cases for which 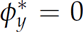. We could not verify this property when non-zero choosiness is selected in both sexes, as we are not aware of the existence of any general method allowing for the assessment of the joint evolutionary stability of several evolving traits.

Second, we analyzed the effect of a change *z* in a given biological or ecological variable on the equilibrium for choosiness in sex *x*, while assuming that other-sex choosiness remains fixed at the equilibrium value reached before the change happens (*i.e*., 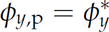). This implies the study of the effect of a change in *z* on the mating rate and/or the expected benefits per mating near 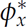 (but not on the expected lifetime because this latter is not related to choosiness). Indeed, at equilibrium we can rewrite eq. 10 as:

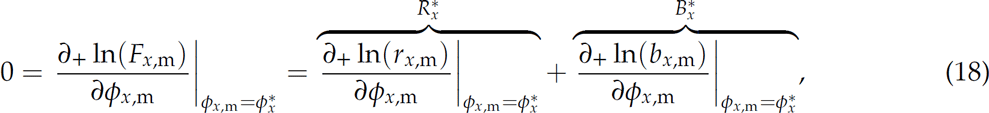

where *∂*_+_ represents the right derivative (*i.e*., we consider the case *ϕ_x_*_,m_ ≥ *ϕ_x_*_,p_ in eqs. 12 & 15, but considering the other case leads to same results as shown in Supplementary Information), 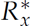 represents the relative change in mating rate in sex *x* at equilibrium and 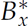 the relative change in expected benefits per mating at equilibrium. Biologically, the value of 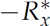 quantifies the decrease in mating rate when choosiness increases, *i.e*., the cost of being choosy. The value of 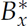 quantifies the increase in expected benefits per mating when choosiness increases, *i.e*., the benefit of being choosy. When *z* influences the mating rate only (hereafter called *z_r_*), we demonstrate in Supplementary Information that (i) the effect of a change in *z_r_* on the evolution of focal-sex choosiness can be deduced from the effect of *z_r_* on 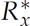 and (ii) this effect can also be deduced from the effect of *z_r_* on the Relative Searching Time (*i.e*., RST: the proportion of lifetime which is devoted to searching for mates):

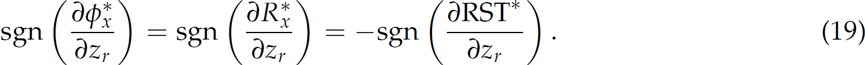

The term 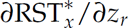 (which is more compactly denoted *∂*RST) corresponds to the sensitivity of RST of the sex *x* with respect to *z_r_*, *i.e*., the variation in the relative searching time caused by the change in *z_r_* while choosiness remains fixed in both sexes.

When *z* influences the expected benefits per mating only (hereafter called *z_b_*), we also demonstrate in Supplementary Information that the effect of a change in *z_b_* on the evolution of focal-sex choosiness can be deduced from the effect of *z_b_* on 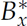:

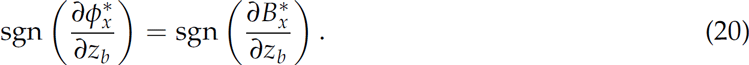

In this situation, we did not find a simple metric such as *∂*RST to summarize the effect of a change in *z_b_*.

Third, we analyzed the effect of a change in *z* on the joint equilibrium for choosiness. Indeed, in the analyzes used to obtain eqs. 19 and 20 we only considered the direct effect of *z* on 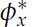 while 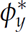 remains fixed, but *z* can also influences 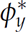, and 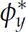 could in turn also influence 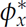. Formally, the total variation of the choosiness in both sexes following a change in *z* is described by the system:

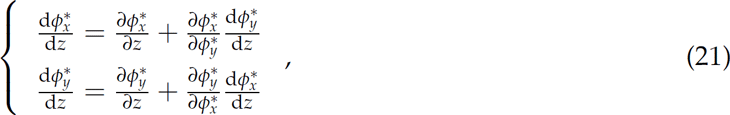

where 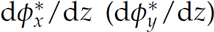 represents the total variation of the choosiness in the sex *x* (*y*) that includes the effect of *z* on the choosiness of both sexes and 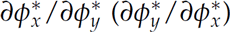 is the variation of 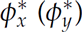 caused by a change in 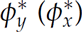 while *z* remains fixed.

We have already described the analysis of 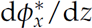 in terms of 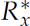 and 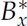, so the same goes for 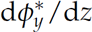 (swapping *x* and *y*). To study 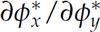 we would similarly consider the changes in 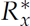 and 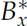 caused by a change in *ϕ_y_*_,p_. However, no more definite analytical result could be obtained for 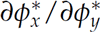 and thus for the overall effect of *z* on the joint equilibrium for choosiness.

### Numerical analysis

Despite the simplicity of the life cycle we consider, some mathematical complexity emerges because of the joint evolution between sexes. As a consequence, some specific results cannot be analytically derived from the equations presented above. We thus complemented the analysis of our model by computing the numerical solution of our analytical equations using the software R (R Core Team, 2015). To minimize the risk of missing exceptions to our main conclusions, we investigated a large number of parameter sets.

To study equilibrium conditions for each choosiness, we considered the 16 possible combinations between 4 different quality distributions for females and males: uniform (*α* = *β* = 1), bell-curve (*α* = *β* = 4), left-skewed (*α* = 4 and *β* = 10) and right-skewed (*α* = 10 and *β* = 4). For each of these 16 cases, we generated two tables of 10^5^ combinations of the other parameters (*γ*, *s_x_*, *s_y_*, *l_x_* and *l_y_*): one for which values for each parameter were randomly drawn from a uniform distribution between 0 and 1 (that we call the “continuous tables”), and the other for which all combinations of values among the following range were considered: 0.001, 0.1, 0.5, 0.6, 0.7, 0.8, 0.9, 0.95, 0.99, 0.999 (that we call the “discrete tables”). In total we therefore analyzed 3.2 *×* 10^6^ (= 16 *×* 10^5^ *×* 2) different parameter sets.

We also used the continuous tables to study the joint evolution of choosiness between sexes. The procedure is described in figure 4.

Finally, we studied the predictive power of *∂*RST numerically. To do so, we first randomly drew 10^6^ pairs of parameters sets differing in the value of only one parameter, from each of the 16 discrete tables. For each pair of parameter sets we computed the partial variation of choosiness, the total variation of choosiness and *∂*RST. Second, we then randomly drew 10^6^ pairs of parameters sets (which could here potentially differ in *γ*, *s_x_*, *s_y_*, *l_x_* and *l_y_*) from the same discrete tables. We computed again the partial variation of choosiness, the total variation of choosiness and *∂*RST for all these pairs. We were therefore able to determine the predictive power of *∂*RST when only one parameter changes, as well as when all parameters are free to change at the same time, using 1.6 *×* 10^7^ (= 16 *×* 10^6^) different parameter sets in each case. The numerical analysis of the predictive power of *∂*RST was not replicated using the continuous table as most of the parameter space sampled in the continuous tables does not lead to situations of mutual mate choice and unilateral choice situations have already been analyzed in Etienne et al. (2014).

## Results

### Scope

We will indicate below whether a given result has been analytically obtained (hereafter labeled as *analytical result*), if it has been obtained for the complete numerical exploration (*numerical result*), or if it has been obtained numerically and correspond to an effect found in only part of the parameter space (*restricted result*). *Numerical results* are consistent across the entire numerical exploration and are likely to be as general as our analytical derivations, that is, true within the scope of the assumptions made in this model. Yet, because this statement cannot be proven without being able to apply a pure analytical approach, we chose to make the distinction between *numerical* and *analytical* results explicit.

### The evolution of mutual mate choice

#### Result 1

There is always one and only one convergence stable (joint) equilibrium for choosiness both in situation of unilateral and mutual mate choice (*numerical result*).

We numerically solved equilibrium eq. 17 for the 3.2 *×* 10^6^ parameter sets and found that there is always one single combination of choosiness that satisfies the equilibrium condition (*numerical result*). For these 3.2 *×* 10^6^ equilibria we found only two outcomes for both convergence and evolutionary stability. First, when the equilibrium is characterized by a null choosiness in at least one sex (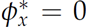 and/or 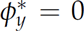), the values of choosiness at equilibrium are the same as in the model of Etienne et al. (2014), in which the choosiness of the non-focal sex was constrained to be null. In this case, the equilibrium is always convergence and evolutionarily stable (*numerical result*). Second, we found parameter settings under which choosiness is non-null at equilibrium in both sexes (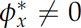 and 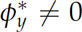), and such equilibria are always jointly convergence stable (*numerical result*). No conclusion could be derived for the evolutionary stability but individual-based simulations for numerous parameters settings suggest that the equilibrium is also evolutionary stable in this case (not shown).

#### Result 2

The fundamental trade-off of mate choice generates a high diversity of combinations of focal-sex and other-sex choosiness at equilibrium (*restricted result*).

Cases of mutual mate choice at equilibrium (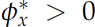 and 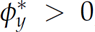) are highly diverse within our numerical exploration, ranging from very low (*e.g*., *ϕ^*^* = 0.01) to very high (*e.g*., *ϕ^*^* = 0.7) choosiness in both sexes, with all possible intermediates (*e.g*., see figure 1).

**Figure 1:**
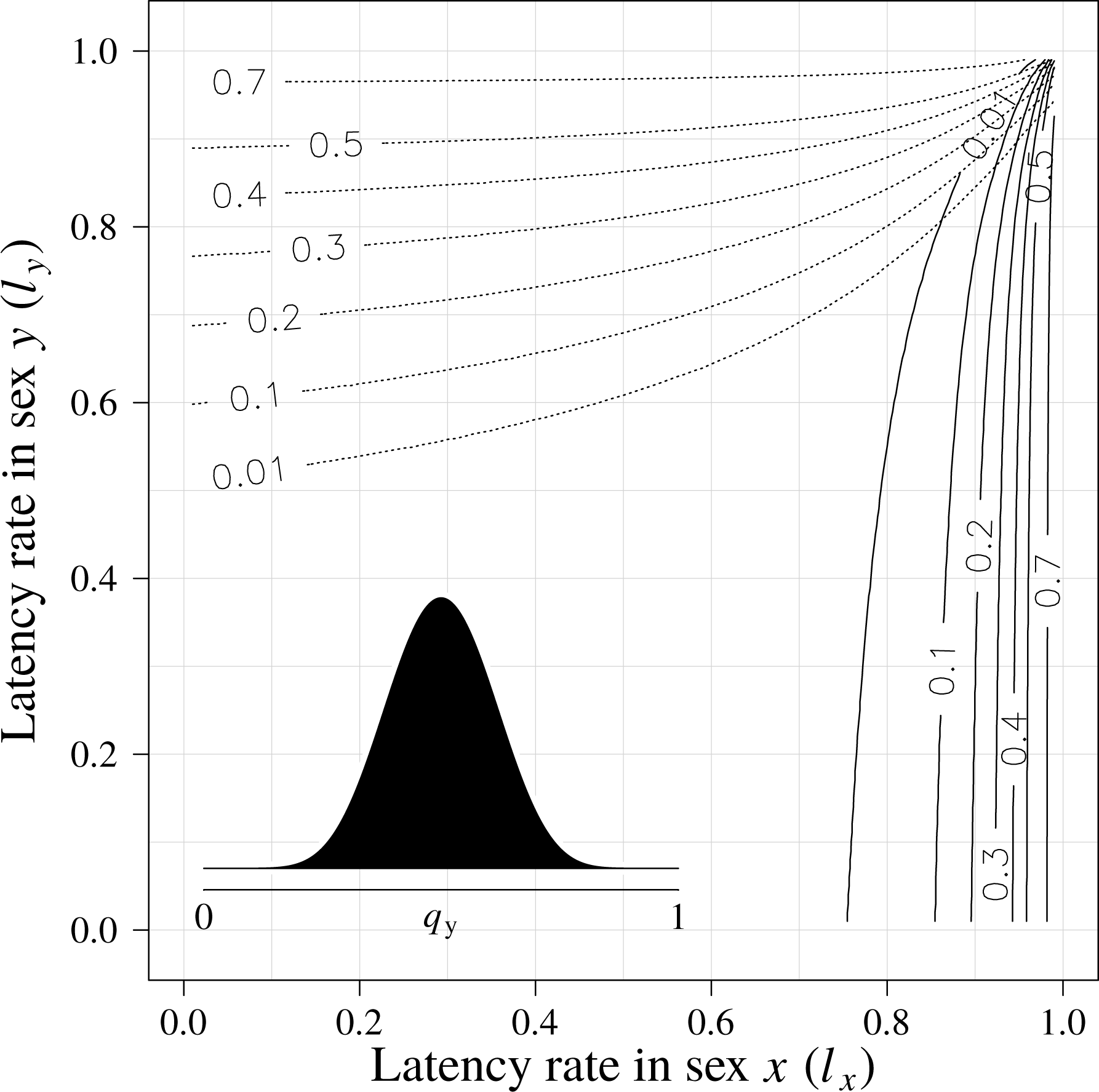
Choosiness at equilibrium in both sexes as a function of latency rates. Contour lines depict the value of choosiness at equilibrium in sex *x* (full lines) and in sex *y* (dotted lines). In this plot the distribution of quality in sex *y* is represented by an insert (*α_y_* = *β_y_* = 4), whereas it is uniform in sex *x* (*α_x_* = *β_x_* = 1), but other distributions are shown in Figure S4. The encounter and survival rates were chosen to favor the evolution of mutual mate choice (*γ* = *s_x_* = *s_y_* = 0.999).

#### Result 3

Within our numerical exploration, mutual mate choice occurs at equilibrium only when both latency and survival rates are high in the two sexes (*numerical result*).

Everything else being equal, the choosier sex is the sex with the (i) higher latency (figure 1 & S4), (ii) higher survival (figure 2 & S5) or (iii) lower variance in quality (figures S4-S6). The evolution of non-null choosiness in a sex requires latency rate in this sex, survival rate in this sex and variance in other-sex quality to be non-null (*numerical result*). However, fulfilling these conditions in both sexes is not sufficient to observe mutual mate choice at equilibrium. Indeed, the latter outcome is obtained only when both latency and survival rates approach 1 in the two sexes (see figures 1-2 & S4-S5). Once this criterion is satisfied, the level of mutual choosiness at equilibrium is influenced by other parameters. In particular, high choosiness in both sexes is favored when encounter rate and/or variance in quality of both sexes is high, and/or mean quality of both sexes is low (figures S4-S6).

**Figure 2:**
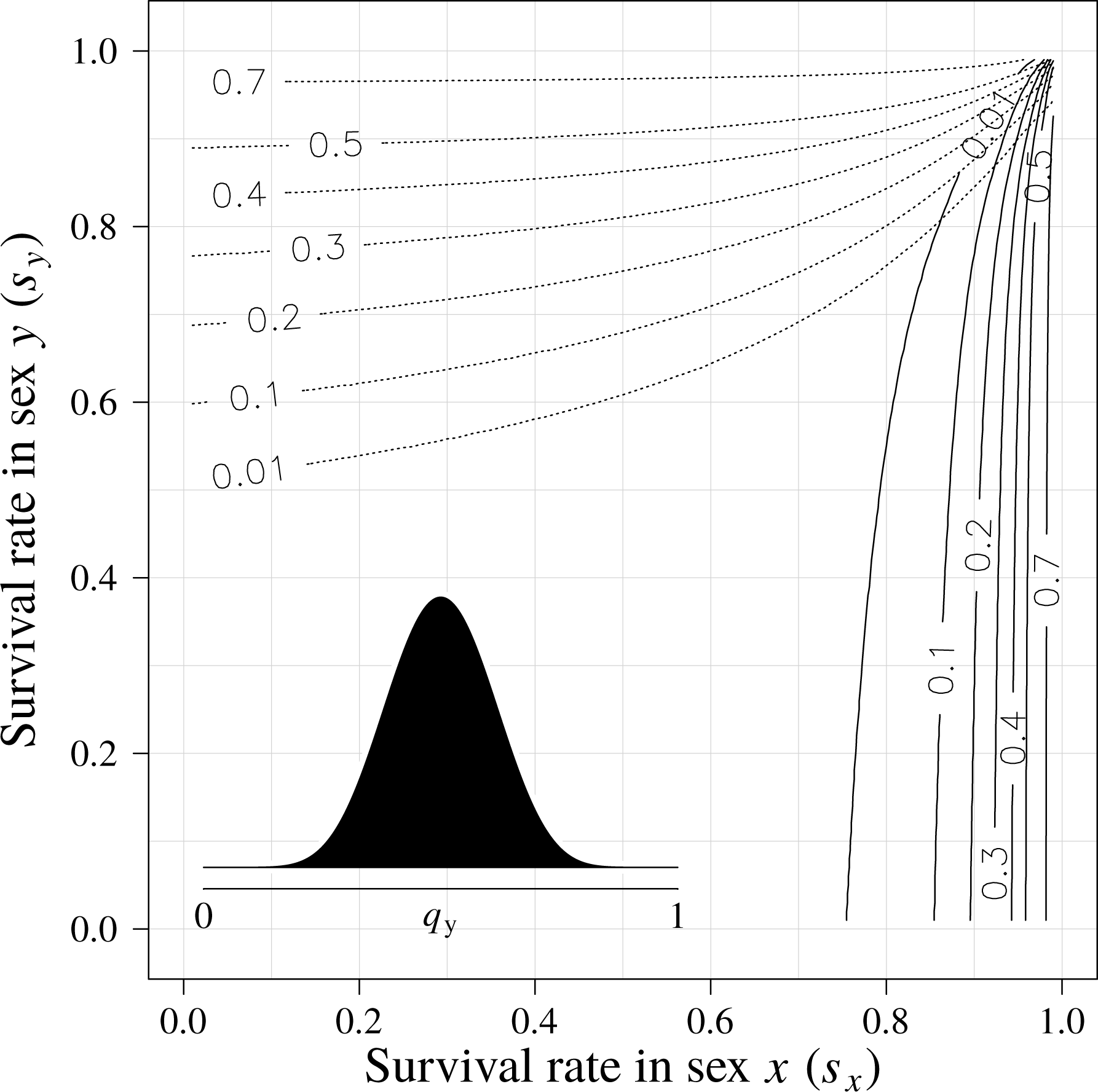
Choosiness at equilibrium in both sexes as a function of survival rates. See legend of figure 1 for details. The encounter and latency rates were chosen to favor the evolution of mutual mate choice (*γ* = *l_x_* = *l_y_* = 0.999).

### The joint evolution of choosiness

#### Result 4

An increase in choosiness in one sex decreases both the cost and the benefit of being choosy in the other sex (*analytical result*).

From the definition of 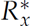 (see eq. 18) and the expression for *r_x_*_,m_ (see eq. 12), the effect of a change in other sex choosiness (*i.e*., *ϕ_y_*_,p_) upon the cost 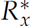 of being choosy is:

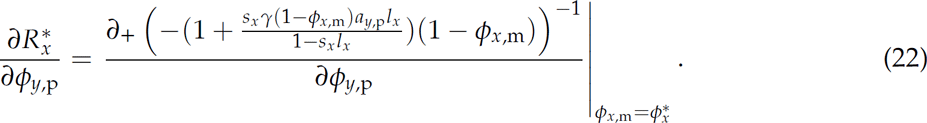

When *ϕ_y_*_,p_ increases, fewer individuals of the focal sex mate, which increases the availability *a_y_*_,p_ of other-sex individuals whose quality is sufficient to mate. Thus the partial derivative of 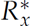 with respect to *ϕ_y_*_,p_ is also positive (*analytical result*). Therefore, an increase in *ϕ_y_*_,p_ selects for higher focal-sex choosiness via its effect on the relative change in mating rate (see eq. 19). Simply put, the increasing availability in the sex *y*, as a consequence of the higher choosiness in this sex, reduces the competition among individuals of sex *x* for the access to other-sex individuals. Thereby the cost of being choosy in sex *x* reduces, which is why *ϕ_y_*_,p_ has, here, a positive effect on 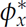.

From the definition of 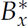 (see eq. 18), the effect of a change in other sex choosiness (*i.e*., *ϕ_y_*_,p_) upon the benefit 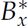 of being choosy can generally be written

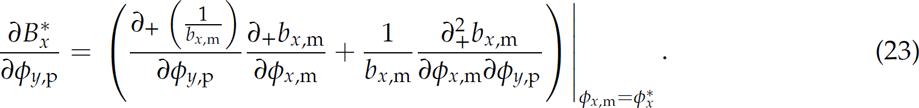

The mixed derivative of *b_x_*_,m_ vanishes (from eq. 15), so that this reduces to

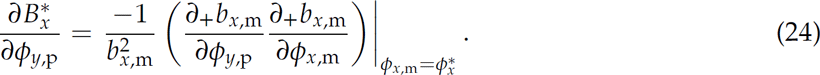

When *ϕ_y_*_,p_ increases, the mean quality of focal-sex individuals whose quality is sufficient to mate increases (see eq. 13), and thus the expected benefits per mating *b_x_*_,m_ increases as well (see eq. 15). Then, *b_x_*_,m_ also increases with *ϕ_x_*_,m_ (see eq. 15). Both derivatives in the right-hand term of the previous equation are thus positive. This implies that the derivative of 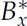 with respect to *ϕ_y_*_,p_ is negative (*analytical result*), and that an increase in *ϕ_y_*_,p_ selects for lower focal-sex choosiness via its effect on the relative change in expected benefits per mating (see eq. 20). To sum up, when choosiness increases in sex *y*, the expected quality of individuals that can qualify as mates increases in sex *x*. This reduces the benefit of being choosy in sex *x*, which implies that *ϕ_y_*_,p_ would thus have a negative effect on 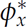.

Had we assumed the reproductive success of a mating pair to be equal to the product of qualities of the two members of the pair (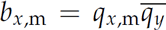), instead of its average (eq. 9), then an increase in other-sex choosiness could have only selected for a higher choosiness in the focal sex (*analytical result*). Indeed, instead of eq. 24, eq. 23 would then lead to:

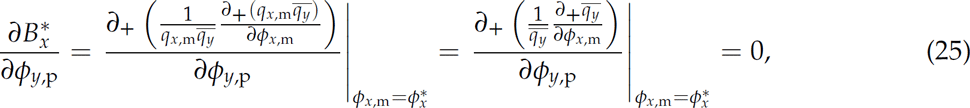

because *q_x_*_,m_ is not a function of *ϕ_x_*_,m_ (see eq. 13) and 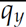 is not a function of *ϕ_y_*_,p_ (see eq. 14). Therefore, the negative effect caused by the influences of *ϕ_y_*_,p_ on the benefit of being choosy vanishes and other-sex choosiness would thus no longer exert a negative effect on focal-sex choosiness. Under this alternative assumption, an increase in *ϕ_y_*_,p_ would thus always lead to an increase in 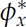 (*analytical result*).

#### Result 5

An increase in choosiness in one sex does not necessarily prevent the evolution of choosiness in the other (*restricted result*).

We have numerically found that when latency rate is low (< 0.7) in both sexes, the negative effect of 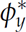 on 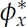 is always larger than its positive effect (*numerical result*, figure 4). However, this result is *restricted* when latency is high in both sexes, which corresponds to cases of mutual mate choice at equilibrium (see figure 1). In this latter situation, parameter values determine which of the two antagonistic effects of 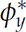 on 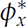 can outweigh the other (figure 4).

### *∂*RST and the effect of a change in mating rate on the evolution of choosiness

#### Result 6

*∂*RST in one sex predicts the evolutionary change in choosiness in this sex so long as the change in mating rate is triggered by variation in a single parameter (*numerical result*).

If a change *z_r_* in a given biological or ecological variable is a function of only one of the parameters which affect the mating rate (*i.e*., *l_x_*, *l_y_*, *s_x_*, *s_y_* or *γ*), then we have found that the partial and total variations of choosiness were always of the same sign for all of the 1.6 *×* 10^7^ combinations of parameters performed (*numerical result* obtained using the discrete tables). This is because in such cases, the partial variation of focal choosiness triggered by *z_r_* outweighs the variation of focal choosiness caused by a change in other-sex choosiness. In these circumstances, computing *∂*RST in a sex is thus sufficient to predict the independent effect of any of these parameters on the evolution of choosiness in this sex, even if this parameter also influences the evolution of choosiness in the other sex. As a consequence, the effects of latency, survival and encounter rates are qualitatively similar between our mutual mate choice model and the one of Etienne et al. (2014) which neglected the effect of a change in other-sex choosiness. Specifically, when latency increases in a sex, *∂*RST is negative for this sex (because lifetime is constant) and positive for the other one (because available mates are rarer), leading to higher and lower choosiness respectively (figure 1). The effect of survival is identical to the effect of latency (figure 2). Indeed, the proportion of the lifetime spent in latency increases with the survival rate in both sexes. This is because when an individual dies, it is always replaced by an available individual, whether the deceased was in latency or not. Finally, when encounter rate increases, *∂*RST of both sexes is negative, which selects for higher choosiness in both sexes (figure 3).

**Figure 3:**
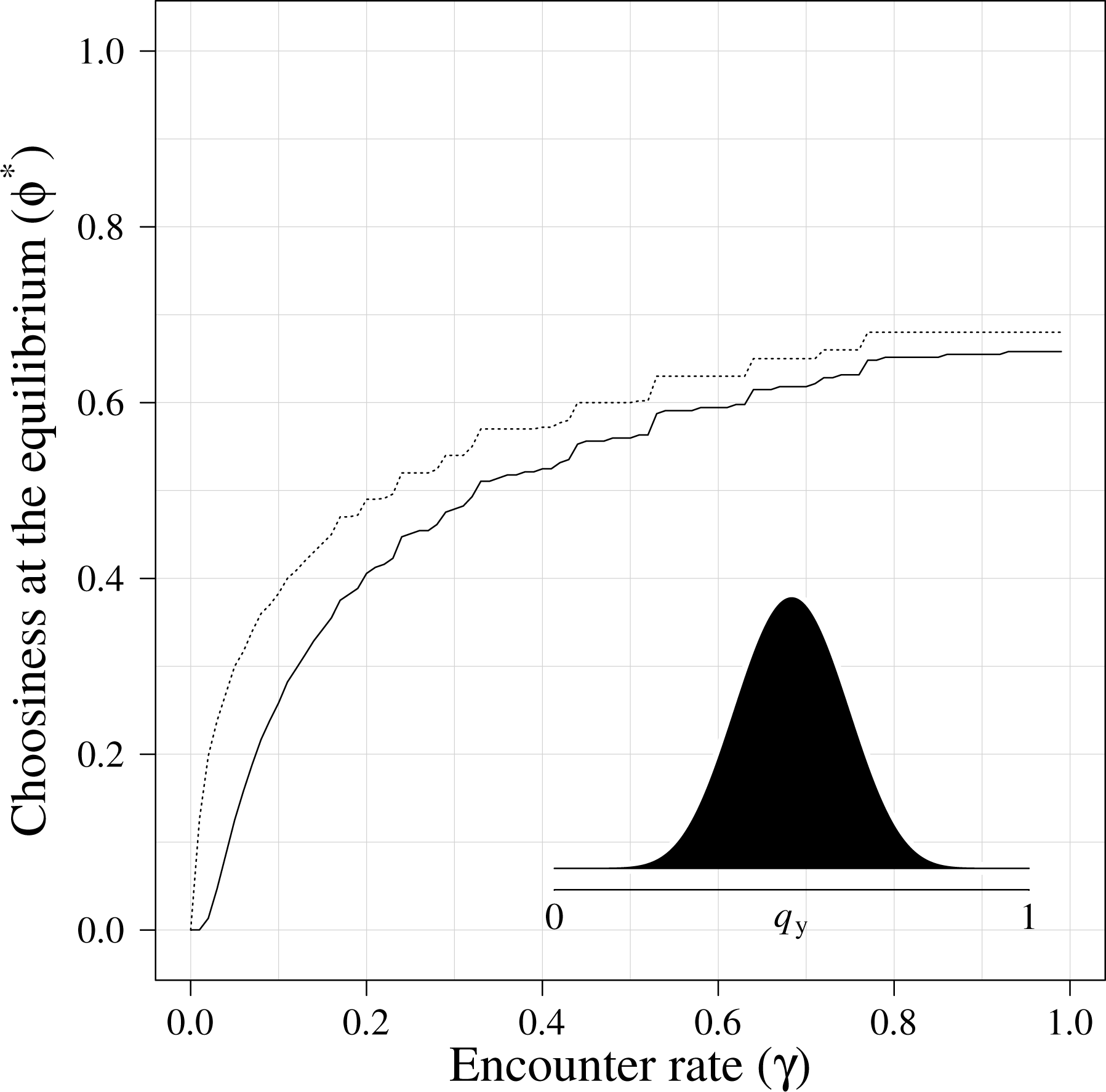
Choosiness at equilibrium in both sexes as a function of encounter rate. See legend of figure 1 for details. The latency and survival rates were chosen to favor the evolution of mutual mate choice (*l_x_* = *l_y_* = *s_x_* = *s_y_* = 0.999). The stepwise aspect of the lines is explained by the use of rounded values for choosiness.

**Figure 4:**
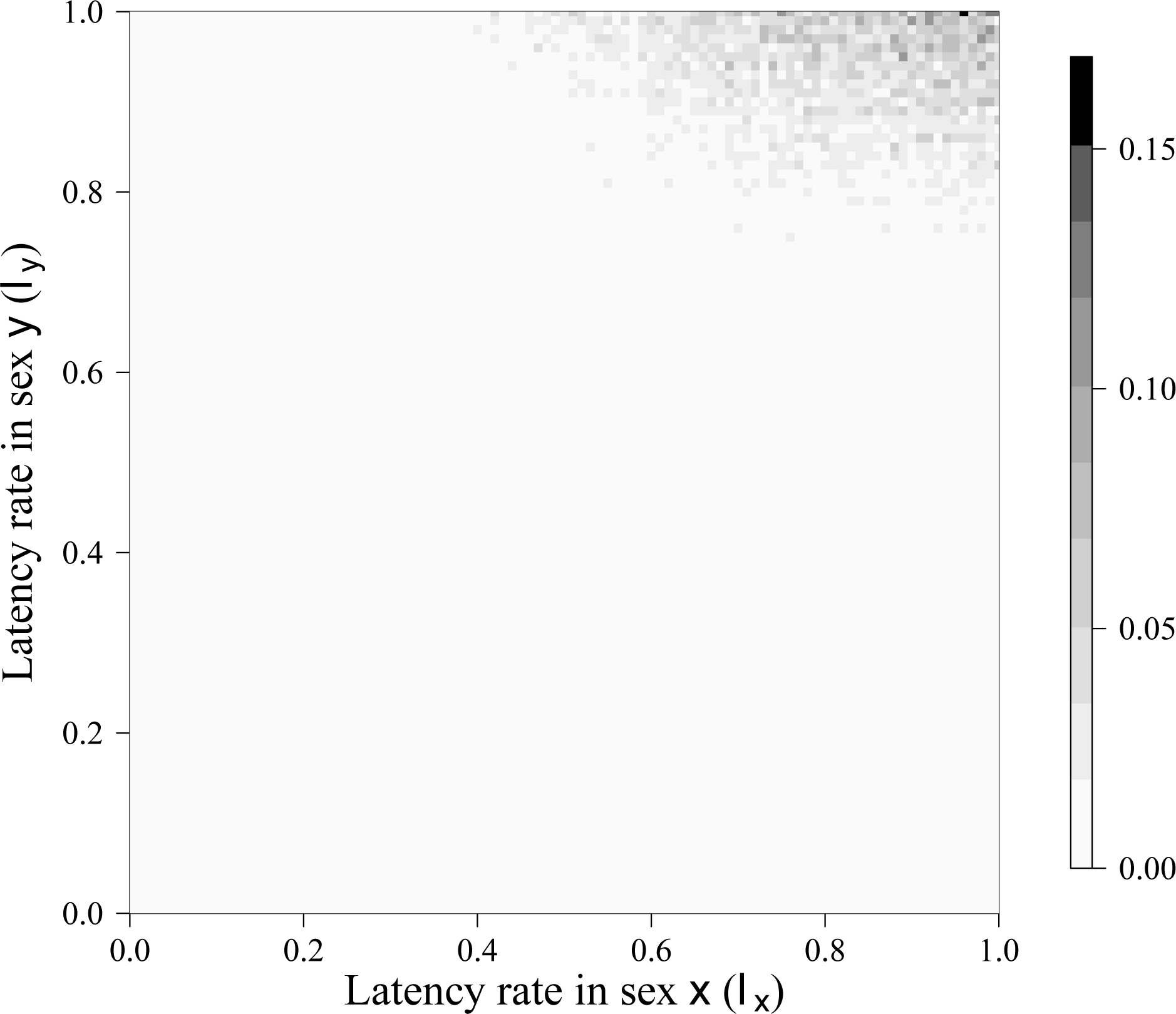
The effect of other-sex choosiness. For each combination of latency rates in sexes *x* and *y*, the color indicates the frequency of cases for which an increase in choosiness in sex *y* has a resulting positive effect on choosiness in sex *x*. This has been obtained by computing the derivative of choosiness in sex *x* at equilibrium (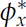) with respect to choosiness in sex *y* at equilibrium (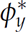) in 1.6 *×* 10^6^ cases exploring the whole range of possible parameter values (using the continuous table, see section *Numerical analysis*). To measure frequencies, the continuous variation in latency was discretized into 101 bins for each axis. The lack of smoothness is explained by the fact that numerical computations are performed for parameters randomly drawn from a uniform distribution. The frequency in each cell of the figure is therefore measured on the variable number of numerical computations (mean ± sd = 156.8 ± 49.5) falling within the corresponding bin for latencies.

#### Result 7

When several parameters vary, the predictive power of *∂*RST is reduced (*restricted result*).

If *z_r_* is a function of more than one parameter, then *∂*RST does not always predict the total variation of choosiness. Indeed, when several parameters affecting the mating rate vary simultaneously, we have numerically found that the variation in choosiness caused by the change in other sex choosiness can outweigh the partial variation of focal choosiness. Cases where *∂*RST loses its predictive power are rare within the parameter space investigated (~0.09%, or 14847 out of the 1.6 *×* 10^7^ combinations of parameters sampled in the discrete tables, see section *Numerical analysis*; figure 5). The cases for which *∂*RST fails to predict the evolutionary change in choosiness are not associated with particular values of the parameters. We found however that *∂*RST can fail when its value is very low (*i.e*., < 0.01) in one sex (this is the case for 8890 out of the 14847 erroneous predictions). It can also fail when both *∂*RST are large. The only structure that we have detected in this latter case is that 84% of erroneous predictions happen when the absolute value of *∂*RST in the focal sex is lower that in the other sex (figure 5).

**Figure 5:**
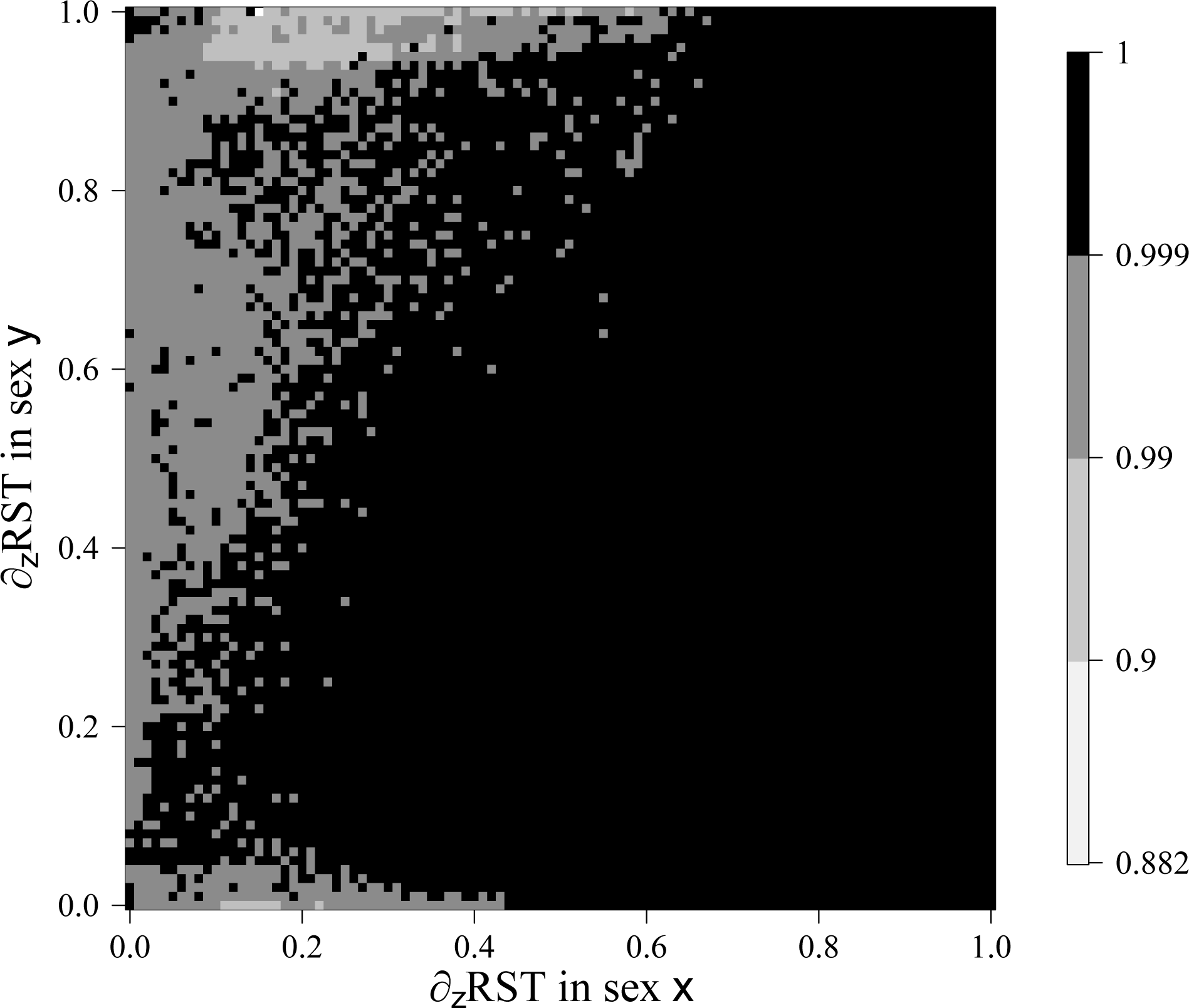
The predictive power of the sensitivity of the relative searching time (*∂*RST). For each combination of the absolute values of *∂*RST in both sexes, the color indicates the frequency of cases for which the sign of *∂*RST in sex *x* correctly predicts the direction of selection of choosiness in this sex. This has been obtained by computing *∂*RST and the total variation of choosiness in both sexes for 1.6 *×* 10^7^ combinations of parameter settings exploring the whole range of possible parameter values (using the discrete tables, see section *Numerical analysis*). Using this approach, the minimal predictive power computed is 88.2%. As in figure 5, *∂*RST was discretized and a variable number of computations falls within each cell, which explains the lack of smoothness (here the number of numerical computations falling within the corresponding bin for *∂*RST is: 1568 ± 3055).

## Discussion

In this article, we have modeled the direct selection of choosiness when mate choice is allowed to evolve in both sexes by considering that mate choice is solely associated with direct benefits in terms of increased mate quality and costs in terms of reduced mating rate. We have neglected all other selection pressures (*e.g*., indirect benefits, energy and predation costs induced by mate search, sexual conflicts) and all other evolutionary forces (*e.g*., drift, migration, recombination). Under these conditions, we derived the complete analytical expression of individual fecundities and obtained most of our results based on the numerical evaluation of our analytical expressions. Opting for a numerical analysis was necessary due to the complexity of our analytical results. This procedure allows for the investigation of the properties of a model under a much larger number of parameter values than when analytical results are lacking (*e.g*., compare our analysis to that of Kokko and Johnstone, 2002). However, a numerical analysis is necessarily less complete than a full analytical study because one cannot *a priori* exclude the possibility that any identified pattern may fail if other parameter values were used. While there is no escape from this general limitation of numerical studies, our analysis explored the entire range of possible values for the life history parameters at a fine scale. For clarity we will therefore label each specific result, as in the previous section, as *analytical*, *numerical* or *restricted* depending on whether it is always true within our set of assumptions, true in our complete numerical exploration, or true for part of the parameter space, respectively.

With these caveats in mind, we have obtained three main results. First, the trade-off between the decrease in mating rate and the increase in benefits per mating (*i.e*., the fundamental trade-off of mate choice) is sufficient to generate the evolution of a high diversity of convergence stable combinations of choosiness between sexes at equilibrium (*Results* 1 & 2 in section *Results*). Within this diversity, mutual mate choice is always characterized by high survival and latency in both sexes but is also influenced by other life history traits (*Result* 3). Second, the evolution of choosiness in a sex can either be promoted or limited by the evolution of choosiness in the other sex (*Results* 4 & 5). Third, *∂*RST (*i.e*., the change in the proportion of lifetime devoted to searching for mates caused only by a variation in any biological or ecological parameter affecting the mating rate of individuals, while choosiness is fixed) correctly predicts the evolution of choosiness in response to a change in mating rate in many but not all cases of mutual mate choice (*Results* 6 &7). We now discuss these results in more detail before examining some key assumptions of our model.

### Life history, through its effect on the fundamental trade-off of mate choice, can select for various convergence stable combinations of choosiness between sexes

Each equilibrium identified during our numerical exploration always corresponds to a single combination of female and male convergence stable choosiness (*Result* 1: *numerical*). Depending on the values of the parameters (encounter rate, sex-specific latency rates, sex-specific survival rates and sex-specific distributions of quality), it is possible to observe a high diversity of values of choosiness at equilibrium in each sex. In particular, all the following combinations can be attained: neither, one, or both sexes are choosy. Cases of mutual mate choice are very diverse, with choosiness ranging from very low (*e.g*., 1% of other-sex individuals are always rejected) to very high values (*e.g*., 70% of other-sex individuals are always rejected) in both sexes. This result leaves open the possibility that direct selection may be sufficient to explain the evolution of mutual mate choice in situations that other studies have interpreted as the result of more complex mechanisms (*e.g*., see Hooper and Miller, 2008; Ihara and Aoki, 1999; Servedio and Lande, 2006; South et al., 2012). In our case direct selection is expressed purely in terms of differential fecundity emerging from differences in the number or in the identity of mates, *i.e*., sexual selection (*sensu* Andersson, 1994, p. 7). Therefore our model challenges the prediction that for mutual choice to evolve one necessary condition is that breeding imposes a large mortality cost on either males or females (Kokko and Johnstone, 2002). Taken together, our model and those of others suggest that there are many paths to mutual mate choice (pre- or post-mating) in nature. In the case of sequential mate choice however, other mechanisms should operate in addition to, and not instead of, the direct sexual selection generated by the fundamental trade-off of mate choice.

In our model, high latency and survival rates in both sexes are necessary for the evolution of mutual mate choice (*Result* 3: *numerical*). Both parameters exert the same effect here because the fraction of the lifetime spent in latency is positively related to both latency and survival rates (see *Result* 6). The latency state in our model can result from any process that prevents individuals from remating instantly, which includes parental investment. Therefore, our findings are consistent with the many empirical studies showing evidence for mutual mate choice in species with biparental care (Amundsen, 2000; Kraaijeveld et al., 2007). Our findings are also consistent with the theoretical studies that showed that a high level of parental investment in both sexes promotes the evolution of mutual choosiness (Crowley et al., 1991; Johnstone et al., 1996; Kokko and Johnstone, 2002; Owens and Thompson, 1994; Parker, 1983). Nonetheless, our definition of latency also encompasses biological situations other than parental investment. Consequently, we also predict mutual mate choice to emerge in organisms that express high latencies for reasons other than high parental investment in both sexes. We therefore predict mutual mate choice to evolve in species in which males suffer high spermatic depletion (because of sperm competition that leads them to produce a high amount of sperm per copulation) and females invest a lot in offspring. This situation may for example explain why in some lekking species such as the great snipe *Gallinago media* (Sæther et al., 2001), or the cichlid fish *Astatotilapia flaviijosephi* (Werner and Lotem, 2003), choice is mutual despite the lack of paternal care. This prediction contrasts with the one made by Kokko and Johnstone (2002) who argued that parental care *per se* and not just mating latency is needed for mutual mate choice to evolve. However, as we shall see later, their assumptions about the mating decision-rule makes the evolution of mutual mate choice more difficult in their case.

The importance of the duration of latency does not preclude other parameters from influencing the level of mutual choosiness (*Result* 3: *numerical*). Indeed, provided that latency and survival rates are high in both sexes, we have obtained predictions similar to those emerging from other theoretical work: high mutual choosiness is favored by a high encounter rate (Crowley et al., 1991; Kokko and Johnstone, 2002), by a high variance in the quality of both sexes (Härdling et al., 2008; Johnstone et al., 1996; Kokko and Johnstone, 2002; Owens and Thompson, 1994 and Parker, 1983) or by low mean quality of both sexes (Gowaty and Hubbell, 2009).

### An increase in choosiness in one sex does not necessarily prevent the evolution of mutual mate choice

In addition to the role played by the aforementioned parameters, we confirmed that the emergence of mutual mate choice can be promoted or constrained by the influence that selection for choosiness in one sex exerts upon selection for choosiness in the other (*Result* 4: *analytical*). Previous work has suggested that the apparent lack of mutual choice in many organisms occurs because an increase in other-sex choosiness may reduce mating opportunities for individuals of the focal sex and would thereby make them less choosy (Kokko and Johnstone, 2002). It is indeed true that if other-sex choosiness does increase, mating opportunities are reduced for low-quality individuals of the focal sex. However, mating opportunities simultaneously increase for high-quality individuals of this sex. Whether this impedes the evolution of mutual mate choice or not is therefore related to the relative extent to which low-quality and high-quality individuals contribute to the gene pool.

In our model, choosiness is expressed as a fixed threshold that is identical for all individuals of a sex. Therefore, we assumed that individuals showing a quality lower than the threshold to be chosen by the other sex do not reproduce at all. As a consequence, only high-quality individuals contribute to the next generation and as such they actually benefit from improved mating opportunities. Formally, when other-sex choosiness increases, the cost of being choosy (*i.e*., the relative decrease in mating rate with choosiness) decreases in the focal sex, which eases the evolution of mutual mate choice in our model. Kokko and Johnstone (2002) assumed a different mating decision-rule. They considered choosiness to be condition-dependent (*i.e*., related to the quality of the individual who chooses), which allows low-quality individuals to pass on their genes to the next generation. Then, the authors observed that the selection pressure caused by the decrease in mating opportunities for low-quality individuals outweighs that caused by the increase in mating opportunities for high-quality individuals, thereby impeding the evolution of mutual mate choice. Therefore, differences between the outcomes of our model and that of Kokko and Johnstone (2002) suggest that the occurrence of mutual mate choice may be strongly influenced by the type of decision-rule individuals use to choose their mates. Empirical knowledge of mating decision rules (*e.g*., Kirkpatrick et al., 2006; Courtiol et al., 2010; Castellano et al., 2012; Reinhold and Schielzeth, 2015) appears therefore crucial for the implementation of realistic models of the evolution of choosiness.

An increase in choosiness in the other sex does not only decrease the cost of being choosy for the focal sex. It also decreases its benefit of being choosy. Indeed, we found that an increase in other-sex choosiness has a positive impact on the mean quality of individuals qualifying as mates in the focal sex, which in turn leads to a reduction of the benefit of being choosy (*i.e*., the relative increase in benefits per mating with choosiness) in this focal sex (*analytical result*). In most of the numerical cases that we have explored, this negative effect on the benefit of being choosy is larger than the cost (*Result* 5: *restricted*), which leads choosiness to decrease in one sex when it increases in the other sex.

Nevertheless, the opposite result can be observed, in particular when latency is high in both sexes, *i.e*., when both sexes are expected to be choosy (*Result* 5: *restricted*). This negative effect of other-sex choosiness on the benefit of being choosy also rests on the questionable assumption that the reproductive success of a mating pair is an additive function of female and male qualities (see eq. 16). Kokko and Johnstone (2002) showed that certain forms of non additive parental care could facilitate the evolution of mate choice. Here, we have shown that this effect is not necessarily limited to care *per se* but can generally emerge from how the fecundity of a pair is determined by the qualities of the two mates. For example, if we consider a multiplicative form for reproductive success instead of an additive one, other-sex choosiness no longer reduces the benefit of being choosy in the focal sex (*Result* 4: *analytical*). Under such an assumption, other-sex choosiness would always promote the evolution of choosiness in the focal sex in our model.

In sum, in terms of joint evolution of choosiness between sexes, the balance between the mechanism selecting for an increase in choosiness and the mechanism selecting against it are strongly dependent on the decision-rule, on how the qualities of mates shape the fecundity of the pair, and of parameter values. Therefore, the only reliable predictions we can propose at this stage are that (i) the evolution of choosiness in one sex can trigger selection pressures both for and against the evolution of choosiness in the other sex, and (ii) the relative effects of these forces are strongly related to biological and ecological factors (*Result* 4: *analytical*).

### The power of *∂*RST as a predictor for the evolution of choosiness must be assessed empirically

We assessed whether one can qualitatively predict an evolutionary change in choosiness triggered by any factor *z_r_* influencing the mating rate of individuals. We found that this is indeed the case, but only under specific conditions. Etienne et al. (2014) showed that one can qualitatively predict an evolutionary change in choosiness triggered by any factor *z_r_* that influences the mating rate of individuals when the other sex is constrained to be indiscriminate. They found that the sign of this change was opposed to the sign of *∂*RST (*i.e*., the variation in the proportion of the lifetime devoted to searching for mates at fixed choosiness). Here, we have assessed the power of this prediction when mate choice is free to evolve in both sexes. When *z_r_* corresponds to a modification of a single parameter in our model, we confirmed this full predictive power of *∂*RST in case of joint evolution of choosiness (*Result* 6: *numerical*). However, when *z_r_* impacts simultaneously on several parameters, this is no longer true as computing *∂*RST in a sex fails to predict the resulting evolutionary change in choosiness in this sex in a few cases of our numerical exploration (*Result* 7: *restricted*). We did not identify any obvious relationship between the predictive power of *∂*RST and the location in the parameter space, but failures occur either when *∂*RST is very small in one sex or when the absolute value of *∂*RST in the focal sex is much larger than that in the other sex (*restricted result*). Importantly, whether the few numerical cases in which *∂*RST fails (on average 9.3 cases per 10,000 trials) represent widespread biological situations or not, is an empirical question – the answer to which shall determine how useful *∂*RST really is.

In cases where *∂*RST accurately predicts the joint evolutionary changes in choosiness, the use of this metric rests on the same three main assumptions as in the model of Etienne et al. (2014): (i) choosiness does not affect survival; (ii) choosiness does not affect the time spent in one latency period; (iii) *z_r_* does not affect the distribution of mate quality, regardless of the form of the latter.

Despite these limitations, we believe that *∂*RST remains superior to all alternative metrics proposed thus far. In particular, Kokko and Monaghan (2001) have clearly demonstrated the limitations of the widely used operational sex ratio (*OSR*). They have suggested using a metric that reflects the cost of breeding (*C*) instead (see also Kokko and Johnstone 2002). While they did so while relaxing our first assumption (*i.e*., choosiness does not affect survival), the predictive power of their metric remains poor: an increase in *C* appears to be a necessary but insufficient condition for the evolution of choosiness in either sex (*e.g*., insufficient when *C* varies from 10^−3^ to 10^−2^ in figure 4 of Kokko and Johnstone 2002; see also Etienne et al. (2014) for an example where *C* produces an erroneous prediction). This weakness emerges from the fact that *C*, as with the *OSR* in many models, is considered as fixed (*i.e*., it depends only on the parameter setting) and does not covary with the evolution of choosiness. Our metric, *∂*RST does not suffer from this limitation (*i.e*., it is internally consistent *sensu* Houston and McNamara, 2005). Therefore *∂*RST captures the complex influence of choosiness on the availability of individuals that qualify as potential partners, which shapes both the benefits and the costs of choice. While additional work may allow for deriving the expression of *∂*RST or a related metric while relaxing assumptions(i) and (ii), a big challenge stems in relaxing the third assumption: as for alternative metrics, the predictive power of *∂*RST rests on the hypothesis that benefits per mating (and thus the distributions of quality) remain unchanged while *z_r_* varies. It would therefore be relevant to identify a predictor that would simultaneously include variations of mating rate and benefits per mating.

In the absence of further developments, we therefore believe that *∂*RST, albeit imperfect, remains the best available predictor of the evolution of choosiness because (i) it holds across a wide range of mating systems, (ii) it encompasses many alternative variables proposed thus far to explain the evolution of choosiness by direct selection (*i.e*., the time invested in breeding, the adult sex-ratio, the operational sex-ratio, and the cost of breeding; see Etienne et al., 2014) and (iii) it can be used empirically to infer qualitative differences in choosiness. We therefore encourage the use of *∂*RST to study the evolution of choosiness in nature both in unilateral and mutual mate choice situations. The guidelines proposed in Etienne et al. (2014) still apply when mate choice is potentially present in both sexes. That is, one should use any proxy that could give an estimation of RST (*e.g*., the time spent sampling mates or courting), and measure this proxy before and after the variable considered has changed (naturally or during the course of an experiment). Then, the difference between the two estimations of RST provide the estimation of *∂*RST. The main empirical constraint is that the first measurement has to be done in a situation in which choosiness is as close as possible to its evolutionary equilibrium in both sexes, and the second before choosiness changes (because of selection or phenotypic plasticity).

Such an experimental protocol aims at predicting the evolution of choosiness in the face of environmental change. If an increase in choice is predicted in one or both sexes, it could also be useful to determine whether the sexual selection predicted to act on mate choice will be strong enough to overcome the influence of other potentially conflicting selection pressures, as well as that of other evolutionary forces. One possibility is to couple the experimental design outlined above with an empirical study in which the environment is maintained constant, the choosiness manipulated, and the mating and reproductive success recorded. Analyzing the outcome of these experiments using the framework of Bateman’s gradients (*e.g*., Anthes et al., 2010; Jones, 2009) should indeed allow the inference of the amount of sexual selection acting on choice in such cases.

Our work should also stimulate empirical perspectives that do not involve *∂*RST. In particular, a precise characterization of the fundamental trade-off of choice in different species would allow the quantification of the direct cost of being choosy, and thereby the assessment of the importance of this trade-off. We are well aware that the empirical assessment of any trade-off is notoriously difficult, however as it has been shown with respect to other questions it is generally worth pursuing (Stearns, 1989). Here, the main difficulty will be – as for the measurement of sexual selection discussed above – to modify the choosiness of individuals without impacting on other parameters influencing the trade-off.

### A critical evaluation of the assumptions of the model

In our model, we have made simplifying assumptions in order to conserve some analytical tractability and thus be able to make general predictions. This naturally raises the question of how robust these predictions are when extended to more realistic and/or specific situations. Due to the complexity of the model, making verbal predictions of the effect of relaxing the key assumptions is highly speculative. Therefore, we encourage theoreticians to build on our formalism to study the effect of some key assumptions that we made for the sake of simplicity. For example, we neglected condition dependence at all levels: choosiness, survival, latency and encounter rate are not influenced by individual quality in our model. This is obviously not realistic (see *e.g*., Cotton et al., 2006) and many other models of the evolution of mutual mate choice have relaxed this hypothesis at some levels (*e.g*., Crowley et al., 1991; Johnstone et al., 1996; Johnstone, 1997; Alpern and Reyniers, 1999; Kokko and Johnstone, 2002). It would be therefore extremely insightful to do the same in our model. We predict that including condition dependence may reduce the predictive power of *∂*RST because this metric does not capture the effect of variables influencing benefits per mating. It may also impede the evolution of mutual mate choice by reducing the decrease that we observed in the cost of being choosy in the other sex. Indeed, in such a case, if assortative mating evolves, low quality individuals should qualify for reproduction, which would produce an effect similar to the one triggered by the mating decision-rule used by Kokko and Johnstone (2002).

A second assumption in our model is that we only consider the evolution of choosiness. However, other traits can evolve jointly with choosiness. Many models focusing on the evolution of mate choice have also focused on how this trait evolves jointly with genetic quality, ornaments (traits indicating the quality of individuals but doing so at a cost), or parental care (for reviews, see Kokko et al. 2006 and Kuijper et al. 2012). While introducing heritable variation in genetic quality in our model would introduce indirect benefits and therefore have a profound impact on the complexity the analyses, ornaments and parental care have been successfully modeled in other work considering only direct selection, even in case of joint evolution between sexes (*e.g*., Kokko and Johnstone, 2002). A natural extension of the present work would therefore be to study how the fundamental trade-off of choice influences the joint evolution between choosiness and these other traits. In the context of this trade-off it would also be interesting to study the joint evolution between choosiness and traits that may mitigate the fundamental trade-off of mate choice. Examples are the evolution of morphological adaptations such as spermathecae that allow female invertebrates to store sperm (Simmons, 2001), or of behaviors such as mate switching during amplexus in male Gammarids (Galipaud et al., 2015). A trivial prediction is that the evolution of such traits should facilitate the evolution of choosiness (in females and males, respectively), but the real question is under which circumstances these adaptations will evolve despite their costs once the benefits of choice are taken into account.

In our model, we have assumed no indirect benefits. This assumption was necessary to study precisely the direct cost that increased choosiness may exert upon the mating rate. Indirect benefits may however occur in nature and strongly influence the joint evolution of mate choice, ornaments and genetic quality (Mead and Arnold, 2004). Studying the role of indirect benefits in mate choice evolution within the framework introduced here may therefore help to study the influence of ecological traits upon the evolution of traits that co-vary genetically with choosiness. It would also help to tackle the controversial topic of the relative role of direct and indirect benefits for mate choice evolution (Kotiaho and Puurtinen, 2007). Moreover, such a model could help to identify the natural conditions for which direct and indirect benefits are aligned (*e.g*., in the case of male choice for sexual swellings in chacma baboons, Huchard et al., 2009) or the conditions for which they are not (*e.g*., in the case of female choice for attractiveness in house crickets, Head et al., 2005).

Finally, we shall discuss one assumption we made that may *a priori* appear limiting but that may not be necessary so: in the real world, latency is not necessarily all-or-nothing in living organisms as assumed in our model, but more likely to vary continuously (one can be more or less available). For example, an individual providing parental care may exhibit an intermediate level of latency since remating is possible, albeit limited, during this period. This seems to contrast with our assumption. However, in our model latency is not all-or-nothing for a group of individuals. This is true in particular for all individuals sharing an allele for choosiness because these individuals will each leave latency at different random times. Hence, selection will be similar at the level of choosiness alleles whether or not latency is, at a given time, all-or-nothing for each individual.

### Conclusion

In this paper, we studied how the choosiness of males and females jointly evolve when selection pressures acting on this trait are only shaped by *the fundamental trade-off of mate choice*: that is the trade-off between the direct benefits individuals gain from choosing their mates and the decrease in mating rate that individuals suffer when they are choosy. We have found that this simple scenario is sufficient to derive several results previously associated with more complex biological assumptions. Contrary to previous claims, we have also revealed that an increase in choosiness in one sex does not necessarily prevent the evolution of mutual mate choice. Indeed, we showed that whether the feedback between the evolution of male and female choosiness promotes or impedes the occurrence of mutual mate choice depends on the life history of individuals (characterized in our model by a survival rate, a latency rate, and an encounter rate, as well as a distribution of the quality of individuals), the decision rule they use for mate choice, and on how the fecundity of a pair is shaped by the quality of both individuals. We have finally demonstrated that *∂*RST, a metric recently proposed in the context of unilateral choice, might also be used to generate global predictions on the evolutionary change in choosiness when mate choice is free to evolve in both sexes.

Our approach reinforces the view that one does not need to enforce any intrinsic difference between the sexes in a model to study “sex roles” (*i.e*., the partition of choosiness and care between females and males). Indeed, we have not constrained life history parameters to particular values according to the sex to which they refer. As such, our model allows the description of the full range of combinations of sex roles regardless of their distribution in nature. However, the same model could be used to tackle questions such as why females are often choosier than males. This could be done by imposing constraints on the parameter values, as others have done (*e.g*., Johnstone et al., 1996).

Our model also highlights the benefits of considering choosiness as a quantitative trait. In particular, our results show that mutual mate choice can be associated with high choosiness in one or both sexes but also with weak choosiness in one or both sexes. The relatively few empirical studies adopting such a quantitative view of mate choice simultaneously in both sexes have already revealed several cases of asymmetric mutual mate choice that were previously documented as unilateral choice (*e.g*., Rowland, 1982; Kraak and Bakker, 1998; Sæther et al., 2001; Werner and Lotem, 2003; Aquiloni and Gherardi, 2008). Pursuing this quantitative approach both theoretically and empirically may lead to greater insights into the frequency of mututal mate choice in nature.

## Acknowledgements

We thank Joel Adamson, Troy Day, Oliver Höner, Olivia Judson, Stephen Proulx, Michel Ray-mond, Robert Schwieger and an anonymous reviewer for helpful comments on the manuscript. Part of the computations were performed on the ISEM computing cluster platform.

